# Efferent Modulation of Spontaneous Lateral Line Activity During and after Zebrafish Motor Commands

**DOI:** 10.1101/769547

**Authors:** Elias T. Lunsford, Dimitri A. Skandalis, James C. Liao

## Abstract

Accurate sensory processing during movement requires the animal to distinguish between external (exafferent) and self-generated (reafferent) stimuli to maintain sensitivity to biologically relevant cues. The lateral line system in fishes is a mechanosensory organ that experiences reafferent sensory feedback via detection of fluid motion relative to the body generated during behaviors such as swimming. For the first time in larval zebrafish (*Danio rerio*), we employed simultaneous recordings of lateral line and motor activity to reveal the activity of afferent neurons arising from endogenous feedback from hindbrain efferent neurons during locomotion. Frequency of spontaneous spiking in posterior lateral line afferent neurons decreased during motor activity and was absent for more than half of swimming trials. Targeted photoablation of efferent neurons abolished the afferent inhibition that was correlated to swimming, indicating that inhibitory efferent neurons are necessary for modulating lateral line sensitivity during locomotion. We monitored calcium activity with Tg(elav13:GCaMP6s) fish and found synchronous activity between putative cholinergic efferent neurons and motor neurons. We examined correlates of motor activity to determine which may best predict the attenuation of afferent activity and therefore what components of the motor signal are translated through the corollary discharge. Swim duration was most strongly correlated to the change in afferent spike frequency. Attenuated spike frequency persisted past the end of the fictive swim bout, suggesting that corollary discharge also affects the glide phase of burst and glide locomotion. The duration of the glide in which spike frequency was attenuated increased with swim duration but decreased with motor frequency. Our results detail a neuromodulatory mechanism in larval zebrafish that adaptively filters self-generated flow stimuli during both the active and passive phases of locomotion.

## Introduction

Sensory systems inform animals about their environment, gathering information essential for directing behavior. Motor behaviors create sensory artifacts that the animal must then disentangle from relevant cues from their environment. In this way, it is critical that sensory systems need to be able to differentiate between external stimuli (exafference) and self-generated stimuli (reafference) to maintain an accurate perception of the sensory landscape (Sperry 1950; Holst and Mittelstaedt 1950). For example, animals that rely on sound-production for communication, such as the cricket, release self-generated acoustic stimuli that can saturate their auditory sensors and cause temporary deafness (Poulet and Hedwig 2006; Crapse and Sommer 2008). A corollary discharge of the sound-producing signal is relayed to the sensory receptor to modulate or interrupt the processing of the reafferent stimulus. By eliminating any ambiguity concerning the origin of a stimulus, animals can better interpret exafferent sensory information by avoiding reafferent sensory overload.

Fishes sense perturbations in the fluid environment through the lateral line, a sensory organ on the body surface that translates fluid motion relative to the body into neural signals essential for navigation (Olszewski, et al. 2012, Suli et al. 2012, Oteiza et al. 2017), predator avoidance, prey capture (McHenry et al. 2009; Stewart et al. 2013), and schooling (Mekdara et al. 2018). However, since water motions are also self-generated by behaviors such as swimming (Palmer et al. 2003; Ayali et al. 2009; Mensinger et al. 2019), respiration (Montgomery et al. 1996; Montgomery and Bodznick 1994; Palmer et al. 2003), and feeding (Palmer et al. 2005), fishes should possess a mechanism to discriminate such self-generated signals from environmental signals.

The origin of this discriminatory capacity is thought to lie in the neural architecture of the lateral line circuit (Russell and Roberts 1972). The neuromast receptors that compose the lateral line are comprised of mechanosensory hair cells innervated by afferent and efferent neurons (Harris et al. 1970; Dow et al. 2018). Afferent neurons (hereafter referred to as afferents) transmit evoked action potentials in response to water motion that deflects the hair cell cilia, resulting in membrane depolarization, rapid glutamate exocytosis, and increased afferent spike rates (Obholzer et al. 2008). Afferents also transmit spontaneous action potentials (spikes) in the absence of mechanical stimuli due to glutamate leakage from the hair cell basal membrane onto their post-synaptic terminals (Keen and Hudspeth 2006; Li et al. 2009; Liao and Haehnel 2012) which is an intrinsic property shown to contribute to maintaining sensitivity (Manley and Robertson 1976, Kiang et al. 1965).

Efferent neuron projections from the brain have long been hypothesized to be a principle mechanism for filtering sensory reafference via efferent copy or corollary discharge (Sperry 1950; Holst and Mittelstaedt 1950). Cholinergic efferent neurons (herein referred to as efferents) projecting from the hindbrain discharge parallel to motor commands that attenuate lateral line activity (Russell 1971; Russell and Roberts 1972; Flock and Russell 1973) as well as activity in homologous octavolateralis systems (i.e. electrosensitive ampullary organ, the inner ear, vestibular system; Bell 1981; Montgomery 1984; Bell 2001, Crapse and Sommer 2008; Weeg et al. 2005; Brichta and Goldberg 2000). The corollary discharge can function to maintain sensitivity to stimulus frequencies during the production of motor commands (Bodznick et al. 1999; Bell 1989; Poulet and Hedwig 2006; Crapse and Sommer 2008). A history of work has been devoted to untangling this critical aspect of sensory systems, often by employing *in vitro* preparations with chemical or electrical stimulation that likely did not match endogenous levels (Russell 1971; Roberts and Russell 1972; Russell and Roberts 1972, Flock and Russell 1973; Montgomery 1984; Tricas and Highstein 1990; Weeg et al. 2005). We expand on this foundational work by investigating the effects of the corollary discharge on lateral line activity *in vivo*, where native levels of feedback are preserved, and defining which parameters of the motor command contribute to lateral line inhibition.

For the first time, we investigate how sensory information from the posterior lateral line of larval zebrafish (*Danio rerio*) is modulated by hindbrain efferents during swimming. Using optogenetic and electrophysiological approaches, we verified the parallel activity of efferents and motor neurons and quantified changes in spontaneous afferent activity during swimming motor commands. To determine the functional role of efferents during swimming, we ablated hindbrain efferents to look for changes in the afferent activity patterns. We next investigated how afferent modulation varies across swim intensities and durations. Lastly, we show that the modulatory effect persists after locomotion and estimated its influence on sensing during the glide phase of intermittent swimming behaviors. Our *in vivo* experimental paradigm is poised to provide a better understanding of sensory feedback during locomotion.

## Methods

### Animals

Experiments were performed on 4-7 day post fertilization (dpf) zebrafish (*Danio rerio*), which were raised in 10% Hank’s solution (137 mM NaCl, 5.4 mM KCl, 0.25 mM Na_2_HPO_4_, 0.44 mM KH_2_PO_4_, 1.3 mM CaCl_2_, 1.0 mM MgSO_4_, 4.2 mM NaHCO_4;_ pH 7.3) at 27°C according to protocols approved by the University of Florida Institutional Animal Care and Use Committee. All larvae displayed a range of swim behaviors (Masino and Fetcho 2005). Blood flow was monitored continuously throughout each experiment as a metric for animal health.

### Calcium imaging

We first verified the cholinergic identity of efferent neurons by backfilling them with tetramethylrhodamine (TRITC, 3 kDa; Molecular Probes, Eugene, OR) in Islet:GFP larvae, as previously (Smith et al. 2014). To image the activity of hindbrain cholinergic efferent neurons, we electroporated TRITC bilaterally into the efferent axons running along lateral line nerve at the site of the cleithrum in 20 Tg(HUC:GCaMP6s) larvae (Zebrafish International Resource Center, Eugene, OR) embedded in agar. Motor neurons were labelled by electroporating TRITC into the axial musculature. Fish were then gently freed from the agar and allowed to recover for 24 hours. Fish were then remounted in agar dorsal surface down and imaged on a Leica SP5 confocal microscope (Leica Microsystems, Wetzlar, Germany). Scanning was performed at 0.39-0.78 Hz, depending on the line speed and scan area. Calcium activity was quantified by a region-of-interest within identified TRITC-labelled cells in ImageJ (v1.48; U. S. National Institutes of Health, Bethesda, MD). We monitored calcium changes in motor neuron activity to indicate swimming. In cases where individual motor neurons did not show calcium changes, we analyze average calcium signal in the spinal cord as a proxy for overall motor activity. Background fluorescence was subtracted by a rolling-ball method implemented in ImageJ, and fluorescence converted to F/F_o_. Baseline fluorescence was manually subtracted using Clampfit software (Molecular Devices, Sunnyvale, CA).

### Electrophysiology

Prior to recordings, larvae were paralyzed by immersion in 10µL of 1mg/mL α-bungarotoxin (Reptile World Serpentarium, St. Cloud, FL) in 10% Hank’s solution. While α-bungarotoxin is an antagonist of the nicotinic acetylcholine receptor (nAChR) α9 subunit in mammalian cochlear hair cells, its effects have been shown to be reversible in *Xenopus* embryos after a 10 minute washout (Elgoyhen et al. 1994). Once paralyzed, larvae were then transferred into extracellular solution (134 mM NaCl, 2.9 mM KCl, 1.2 mM MgCl_2_, 2.1 mM CaCl_2_, 10 mM glucose, 10 mM HEPES buffer; pH 7.8, adjusted with NaOH) and pinned with etched tungsten pins through their dorsal notochord into a Sylgard-bottom dish.

Multi-unit extracellular recordings of the posterior lateral line afferent ganglion were made in wild-type ABTü fish (n = 29). Electrodes (30µm tip diameter) were pulled from borosilicate glass (model G150F-3, inner diameter: 0.86, outer diameter: 1.50; Warner Instruments, Hamden, CT) on a model P-97 Flaming/Brown micropipette puller (Sutter Instruments, Novato, CA). The recording pipette was filled with extracellular solution and brought into contact with afferent somata by applying −50 mmHg pressure (pneumatic transducer tester, model DPM1B, Fluke Biomedical Instruments, Everett, WA). Negative pressure was released to atmospheric (0 mmHg) once a stable recording was achieved. This approach allowed us to record the activity of multiple posterior lateral line afferents and increase our chances of observing any effect during motor activity.

Along with afferent recordings, simultaneous extracellular ventral root (VR) recordings were made through the skin (Masino and Fetcho 2005) between myotomes 6-10, where myotome 1 is the anterior-most muscle block. VR pipettes were beveled and polished to a >60 µm tip diameter using a MF-830 Microforge (Narishige International, Amityville, NY). VR pipettes were held at −100 mmHg during recordings. Spontaneous swimming activity was recorded simultaneously with evoked swimming commands, which were initiated by light or an electrical pulse (60 V for 20 ms, model DS2A – mk. II Constant Voltage Isolated Stimulator, Digitimer Ltd., Welwyn Garde, UK). All recordings were sampled at 20 kHz and amplified with a gain of 1000 in Axoclamp 700B, digitized with Digidata 1440A and saved in pClamp10 (Molecular Devices).

All recordings were analyzed in Matlab (vR2016b) using custom written scripts. Both spontaneous afferent spikes and swimming motor activity identified using a combination of spike parameters such as threshold, minimum duration (0.01 ms), and minimum inter-spike interval (ISI; 0.001 ms). Motor bursts within a swim bout were then defined by any activity that had a minimum of two spikes within 0.1 ms of each other and lasted a minimum of 5 ms. All swim bouts that were selected for analysis had a minimum of three bursts with a maximum inter-burst interval of 200 ms. We observed 2,336 total swim bouts, of which 45.8% were preceded by intervals of inactivity in the afferent neurons. These periods of inactivity made it challenging to interpret changes in afferent activity, so we restricted the dataset to only include swim bouts that were preceded by a minimum of one afferent spike (n = 1,263 swim bouts). This period of monitored afferent activity was set to match the time interval of the subsequent swim bout. For the rest of the paper, this period is termed “pre-swim”, with the same time period following a swim bout termed “post-swim”. The interval between the end of a swim bout and the first afferent spike was termed the “refractory period”.

### Efferent Neuron Ablations

ABTü wild-type fish (n = 8) were mounted on their sides in 1.6% low-melting point agarose, and covered in 0.02% MS-222 in Hank’s solution. The somata of efferent neurons were backfilled as previously by TRITC (3 kDa) or fluorescein isothiocyanate (FITC, 3-5 kDa, Sigma-Aldrich, Darmstadt, Germany), and allowed to recover for 2-4 hours. The near-ultraviolet laser was focused at a depth corresponding to the maximum intensity of each neuron’s soma, to ensure we were targeting its center on a Leica SP5 TSM laser scanning confocal microscope. Labelled efferents were imaged at 532 nm, and we applied the FRAP Wizard tool in Leica application software to target individual cells. We ablated target cells with a 30 s exposure to the near-ultraviolet laser line (458 nm), and successful targeting was typically confirmed by quenching of the backfilled dye. This method has been successfully applied and validated in similar systems (Soustelle et al. 2008; Liu and Fetcho 1999). Fish were then freed from agar and allowed to swim freely and recover overnight. Electrophysiological recordings were performed to simultaneously record afferent activity and motor activity in ablated animals. If swimming has no impact on spike rate, then the null hypothesis is that the slope of the line of best fit forced through zero will not be significantly different from unity. Conversely, a departure from this slope indicates that swimming has a significant impact on spike rates: a higher slope corresponds to increased spike rates, and lower slope corresponds to suppressed spike rates.

### Behavioral Test

ABTü wild-type fish (4-7 dpf, n = 50) were housed in a 400 mL beaker in 200 mL of Hank’s solution. Intermittent swimming behavior of larvae were recorded from above using a high-speed video camera (200 frames per second, 512×512 pixel resolution, 1600 ms exposure time, Phantom Miro M310, Vision Research Inc., Wayne, NJ, USA) and bottom lit by an LED array (Ikan Corp., Houston, TX) through a diffuser. To prevent microcurrents arising from thermal gradients caused by the LED lighting (which we found to prevent accurate measurements of the glide phase), we introduced a fan to circulate air and offset the heat. Video capture was triggered to record the swim and the glide of individual larvae from initiation of the tail-beat until lateral movement ceased. Videos (n = 112) were then processed using custom written Matlab programs to track the body kinematics of larvae, frame-by-frame, and quantify tail-beat frequency and swim duration. The glide period was defined as the time from the end of the tail-beat to the end of passive lateral movement.

### Analysis

All statistics were performed using custom written models in the R language (R development core team, vR2016b) using packages car, visreg, reshape2, plyr, dplyr, ggplot2, gridExtra, minpack.lm, nlstools, investr, and cowplot (Fox and Sanford 2011; Breheny and Burchett 2017; Wickham, 2007, 2011, 2016; Wickham et al. 2019; Augule 2017; Elzhov et al. 2016; Baty et al. 2015; Wilke 2019). We were able to measure changes in calcium activity in 9 out of 10 efferents, and these neurons were subsequently analyzed. We analyzed the correlation between the time courses of efferent neurons and motor neuron calcium activity using cross-correlations of the respective time series. Pre-swim, swim, and post-swim spike rate were calculated by taking the number of spikes within the respective period over its duration. Swim frequency was calculated by taking the number of bursts within a swim bout over the duration of the swim bout, and duty cycle was calculated by taking the sum of the swim burst durations over the swim bout’s total duration. Relative spike rate was calculated by taking the swim spike rate over the pre-swim spike rate. All variables were averaged by individual (n = 29). The precision of estimates for each individual is partly a function of the number of swims, so we analyzed variable relationships using weighted regressions, with individual weights equal to the square root of the number of swims. We log transformed variables in which the mean and the variance were correlated. To quantify the inhibition of the afferent spike frequency during swimming we tested for a significant difference in afferent spike frequency during swimming as compared to non-swimming periods using a paired sample student’s t-test.

Differences in afferent spike rates across the various periods of interest (pre-swim, swim, and post-swim) were tested by two-way analysis of variance (ANOVA) followed by Tukey’s post-hoc test to detect significant differences in spike rates between swim periods or treatments. Linear models were used to detect relationships between spike rate during swimming and other independent variables (e.g. spike rate pre-swim and motor parameters). To quantitatively define the refractory period, we restricted the data to individuals that had a minimum of 20 swims and sampled spike times within 500 ms following the termination of the swim. We used a modified logistic function to model asymmetrical data from parameters of post-swim spike rates to predict the duration of the refractory period, according to the equation,

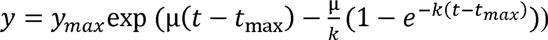

where the response *y*. at time *t* is due to exponential growth with constant *k* until reaching a maximum *y*_max_ at time *t*_max_, after which it grows as μ (Werker and Jaggard 1997). The model was bootstrap simulated for 100 iterations.

We used behavioral data to quantify the relationships between tail-beat frequency and swim duration to glide duration, as described above. We then calculated the expected refractory period duration of the behavioral motor parameters from the line equations produced by linear models of the ventral root motor parameters. With the expected refractory period and respective glide duration we calculated the proportion of time the refractory period overlapped the glide phase and quantified the relationship between tail-beat frequency and swim duration to that proportion. Data is shown throughout the manuscript as mean ± standard error. Statistical significance is reported at α = 0.05.

## Results

We first quantified the strength of the association between efferent neuron activity and motor activity. Our optogenetic approach confirmed that efferent neurons on both left and right sides of the hindbrain are active during swimming. Cross-correlation coefficients of efferent neuron and motor activity time profiles ranged 0.44 to 0.95, with a mean of 0.66. In one example of an individual with four neurons labelled on each side, the correlation coefficient was 0.89 (Figure 1). We only observed strong calcium activity in efferent neurons during swimming. Therefore, we performed ventral motor root recordings to signal the timing of efferent activity and to dissect the components of the motor signal that are transmitted to the lateral line during the corollary discharge.

**Figure 1.**
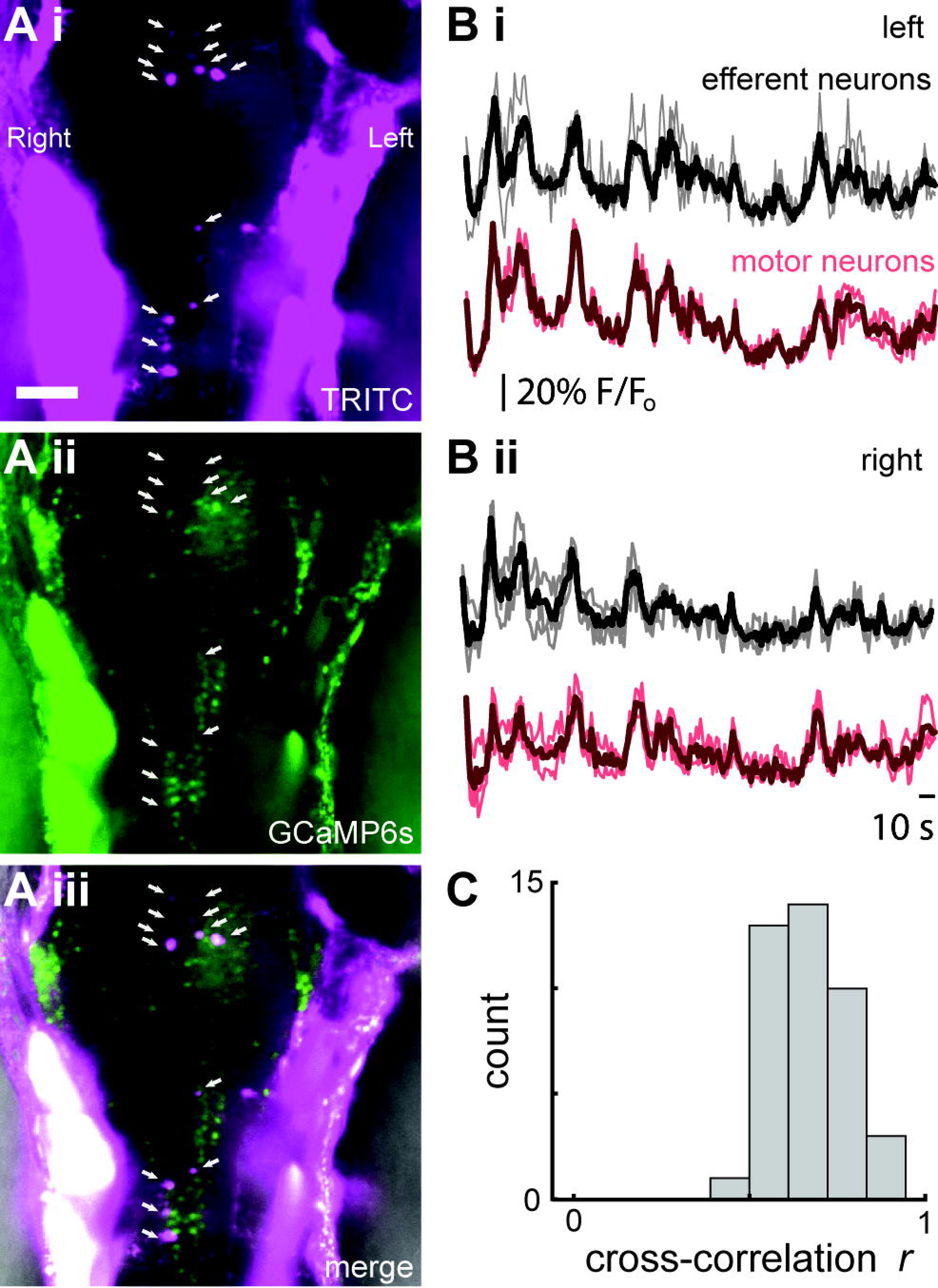
Optogenetics reveals synchronous activity of cholinergic efferent neurons and motor neurons in the hindbrain. **Ai-iii.** Efferent cell bodies (arrows) were positively identified and labeled by backfilling the lateral line neuromasts with TRITC in TG(HUC:GCaMP6s fish. Scale bar is 50 µm. Labelled cells from which calcium activity was optically recorded are designated with an arrow. **B.** Example from one representative animal showing changes in calcium activity as indicated by fluorescence profiles in both efferent and motor neurons. Data are shown for both left (i) and right (ii) sides of the body, where dark traces are average values across all cells and light traces denote individual neurons. Scale bar is 10 seconds. **C.** Cross-correlation coefficients between all efferent and motor neurons.

Spontaneous afferent activity was reduced during swimming, but is not fully inhibited in all cases (Figure 2). The distribution of spike rates was shifted towards smaller values which results in a significant decrease in afferent spike rate from when the larvae were inactive (5.90 ± 0.1 Hz) compared to when they were swimming (3.81 ± 0.2 Hz; p < 0.001, t = 9.88, df = 3,236; Figure 2 D). We examined spike rates during swimming relative to the pre-swim period to examine patterns of relative inhibition. In a majority of swim bouts, there was a complete absence of afferent activity (relative spike rate = 0, n = 620/1,263, 50.7%), but there were also instances of partial decrease (relative spike rate between 0 and 1, n = 267/1,263, 21.1%), no decrease (relative spike rate = 1, n = 240/1,263, 19.0%), and an increase in spike rate (relative spike rate > 1, n = 116/1,263, 9.2%; Figure 3 A, B). Discrete peaks in relative spike rates arise due to the integer number of spikes during swimming, often due to multiples of one.

**Figure 2.**
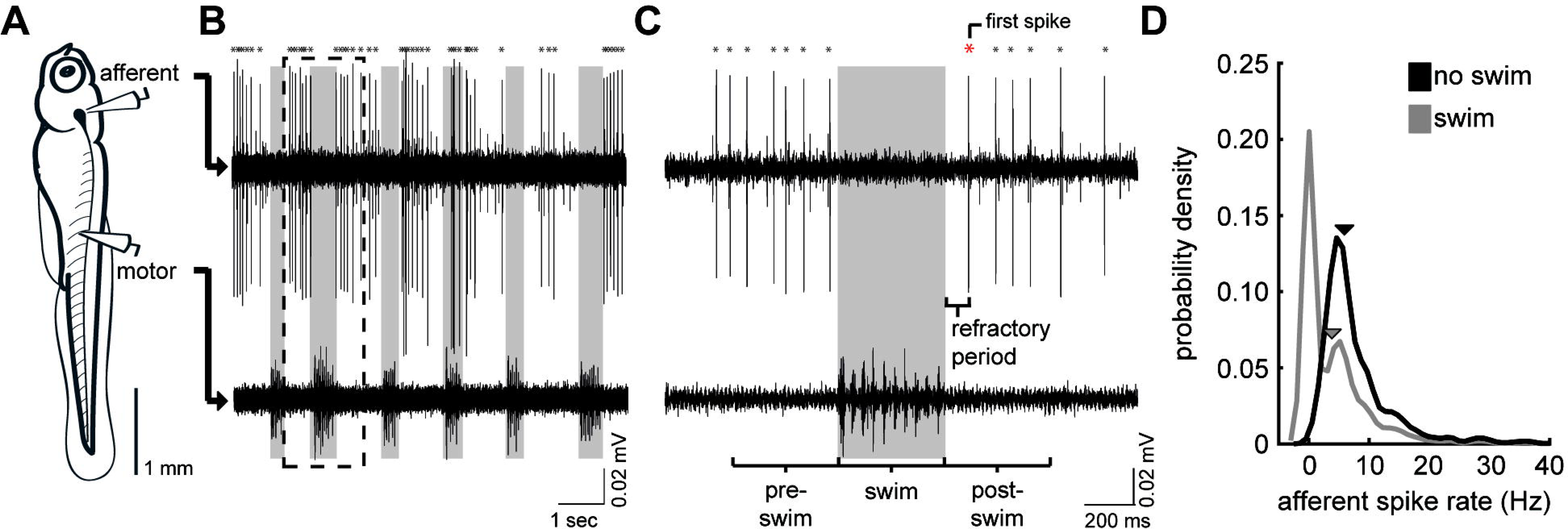
Swimming inhibits spontaneous lateral line afferent activity. **A.** Simultaneous recordings from afferent neurons from the posterior lateral line afferent ganglion and ventral root recording of motor neurons along the body were made in 29 paralyzed larval zebrafish between 4-7 dpf. **B.** At the onset of each fictive swimming bout, the spontaneous spike rate of afferent neurons decreases. **C.** Dashed box in B shows that in this example spontaneous afferent activity is completely inhibited (gray box). We used three intervals of interest relative to the swim duration to assess changes in spike rate, pre-swim, swim, and post-swim. We also examined the interval between the termination of motor activity and initiation of the first spike after the swim which we define as the refractory period. Asterisks at the top of the panel denote afferent spike that exceed the minimum threshold for our spike detection algorithm and the red asterisk indicates the first spike after the swim and termination of the refractory period. **C.** Kernel density estimate of spike rates during swimming versus non-swimming with arrows indicating the mean (n = 1,263).

**Figure 3.**
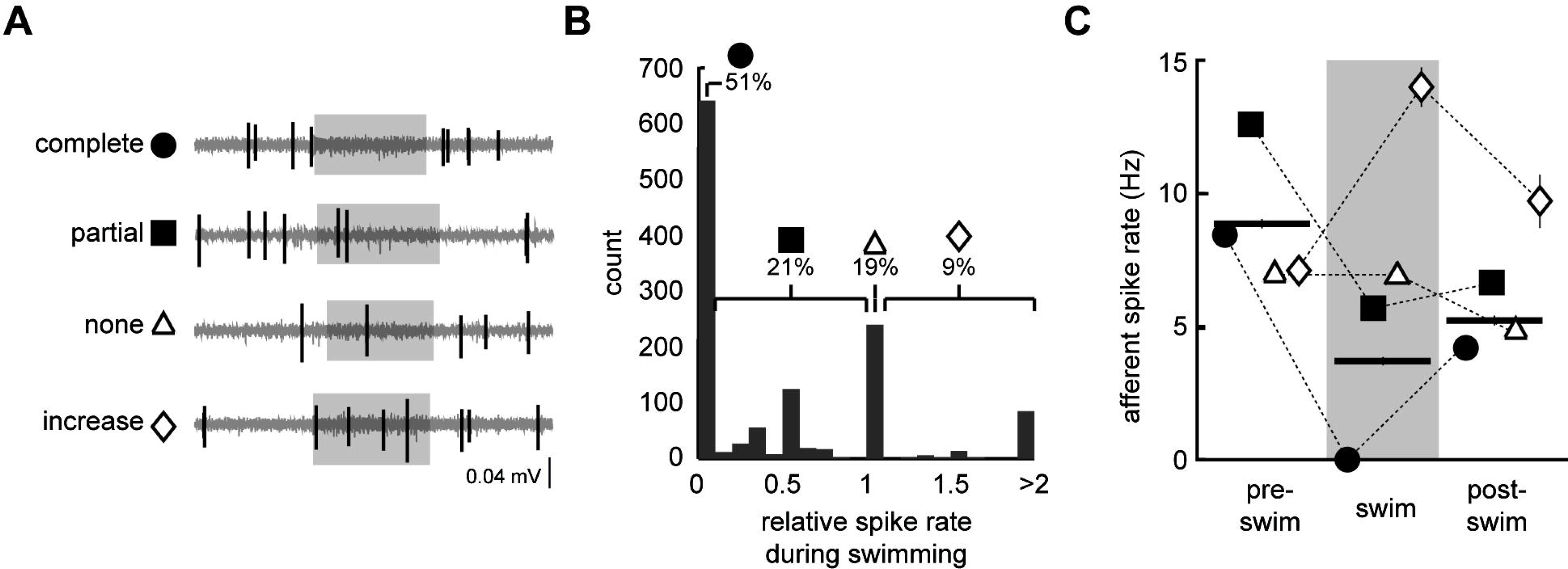
Quantification of spontaneous afferent spike rate during fictive swimming. **A.** Example from a representative individual illustrating different levels of spontaneous afferent activity during swimming; complete decrease (filled circle), partial decrease (filled square), no change (open triangle), and an increase (open diamond) in afferent spike rate. Note that spontaneous spike rate was determined to be able to assess these definitions (see Methods). **B.** Histogram of spike rates during swimming relative to pre-swim spike rate, showing that over half of the trials consisted of completely inhibited afferent activity. **C.** Afferent spike rates during pre-swim, swim, and post-swim (horizontal bars) during instances of complete decrease, partial decrease, no change, and increased spike rates. Error bars represent ± SEM.

In order to more closely probe the dynamics of afferent activity, we examined how spike rate varies before, during, and after the swim. For this analysis we only examined those swims with spikes that occurred within the pre-swim interval. Swimming spike rates were lower than the immediate pre-swim period (8.94 ± 0.2 Hz, relative decrease 57%) and the post-swim period (5.34 ± 0.2 Hz, relative decrease 40%). Post-swim spike rate was, itself lower than the pre-swim spike rate (Tukey post-hoc tests across groups, p < 0.001). Spike rates preceding a complete decrease (8.45 ± 0.2 Hz), no decrease (6.94 ± 0.3 Hz), or an increase in spike rate (7.15 ± 0.5 Hz) were significantly lower than spike rates preceding partial inhibition (12.64 ± 0.4 Hz; F_3,1262_ = 50.03, p < 0.001; Figure 3 C).

We expected that ablating the efferent neurons would diminish or abolish the swim-associated inhibition of spike rate. Following efferent ablation, we observed some decrease in afferent spike rates but to a lesser extent than in non-ablated fish which was not statistically discriminated from pre-swim spike rates of either group (Tukey post-hoc test; ablated, pre-swim: 8.92 ± 0.268 Hz; ablated, swimming: 5.65 ± 0.3 Hz; Figure 4 B). Alternatively, we can examine the correleation between non-swimming and swimming spike rates for evidence of the effects of suppression. If swimming has no impact on spike rates, then the line of best fit will be unity because on average, swim intervals are equally likely to be higher or lower than non-swim intervals. Average spike rates during and prior to a swim were positively correlated in control fish (r^2^ = 0.61, F_1,35_ = 54.48, p < 0.001), and the slope of the line indicates a fractional suppression of 56% that is significantly less than unity (slope 0.44, CI: 0.32-0.57; Figure 4 C). Conversely, in ablated fish, spike rates during swimming are not distinguishable from unity (slope 1.12, CI = 0.78-1.46), indicating that spike rates during swimming intervals differs from non-swimming interval no differently than chance.

**Figure 4.**
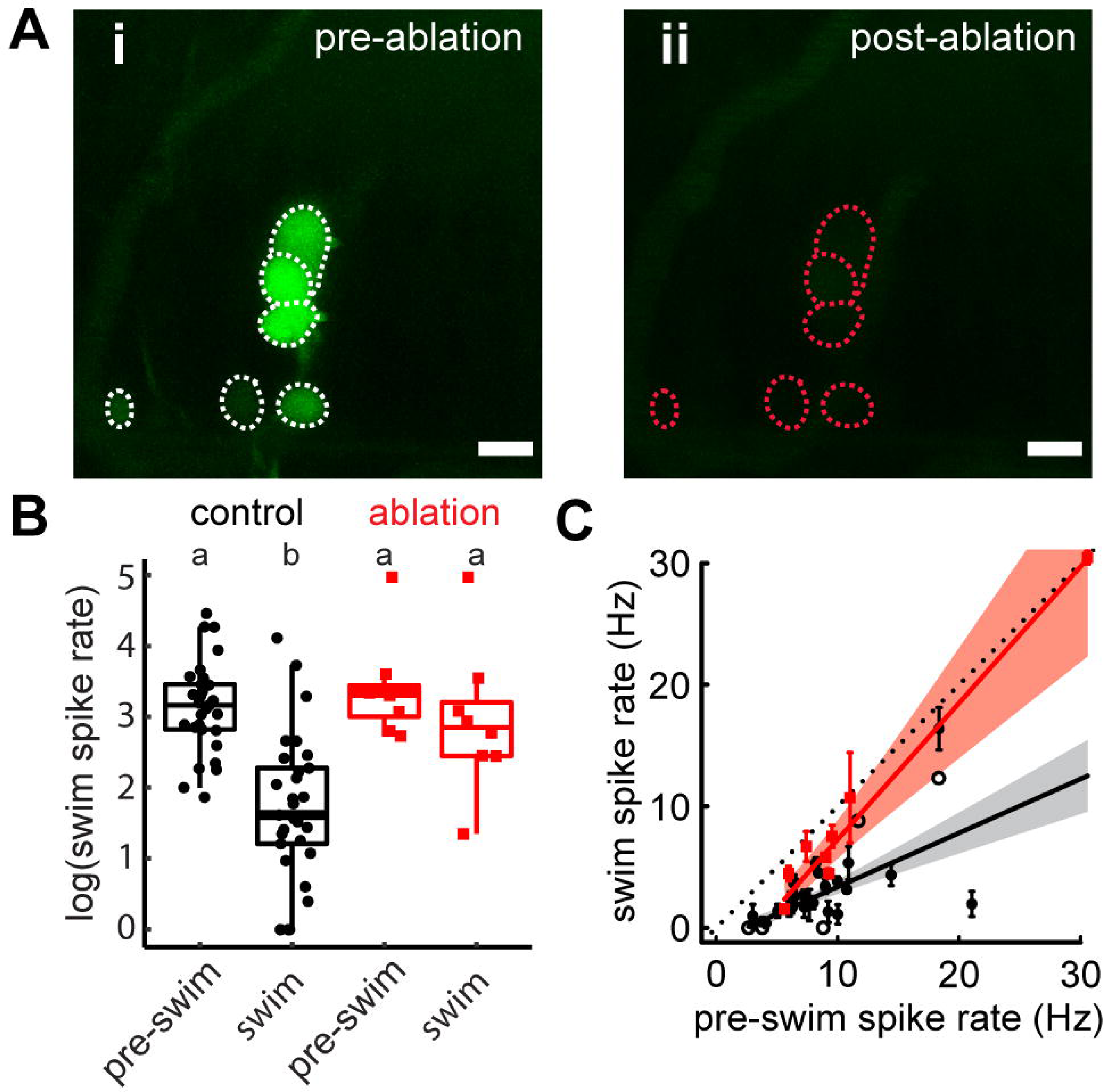
Cholinergic efferent neurons are necessary for afferent inhibition during swimming. **A.** Efferent cell bodies were identified by backfilling fluorescein through the lateral line (i) and ablated with a 30 s UV pulse on a Leica confocal microscope (ii). Scale bar represents 10 µm. **B.** Individuals with ablated efferents displayed reduced spike rate during swimming compared to control individuals. **C.** The line of best fit of spike rates before compared to during the swim significantly excludes unity in control fish but not ablated fish, implying spike rate suppression in the former but not the latter. Dashed line indicates the line of unity, corresponding to no average difference of spike rate during swimming. All error bars represent ± SEM.

We sought to determine what aspects of the age dependent motor pattern may contribute to the corollary discharge transmitted to the lateral line. Across ages, average swim frequency, duty cycle, and swim duration were 27.01 ± 6.0 Hz, 0.101 ± 0.002, and 280 ± 6.0 ms respectively. Swim frequency and duty cycle both increased with age, while swim duration decreased with age (r^2^ = 0.378, F_2,27_ = 16.41, p < 0.001; r^2^ = 0.437, F_2,27_ = 20.93, p < 0.001; r^2^ =0.166, F_2,27_ = 5.377, p = 0.028, respectively; Figure 5 A-C). Relative changes in spike rate during swimming did not change with age (r^2^ = 0.047, F_2,27_ = 1.331, p = 0.259). Swim frequency and duty cycle were not correlated to changes in spike rate (r^2^ = 0.099, F_2,26_ = 1.431, p = 0.231and r^2^ = 0.047, F_2,26_ = 0.645, p = 0.932, respectively; Figure 6 A, B), but the spike rate was predictably lower for longer swim durations (r^2^ = 0.186, F_2,26_ = 2.971, p = 0.045; Figure 6 C). Mixed effect models revealed no interaction between frequency, duty cycle, and duration with individuals.

**Figure 5.**
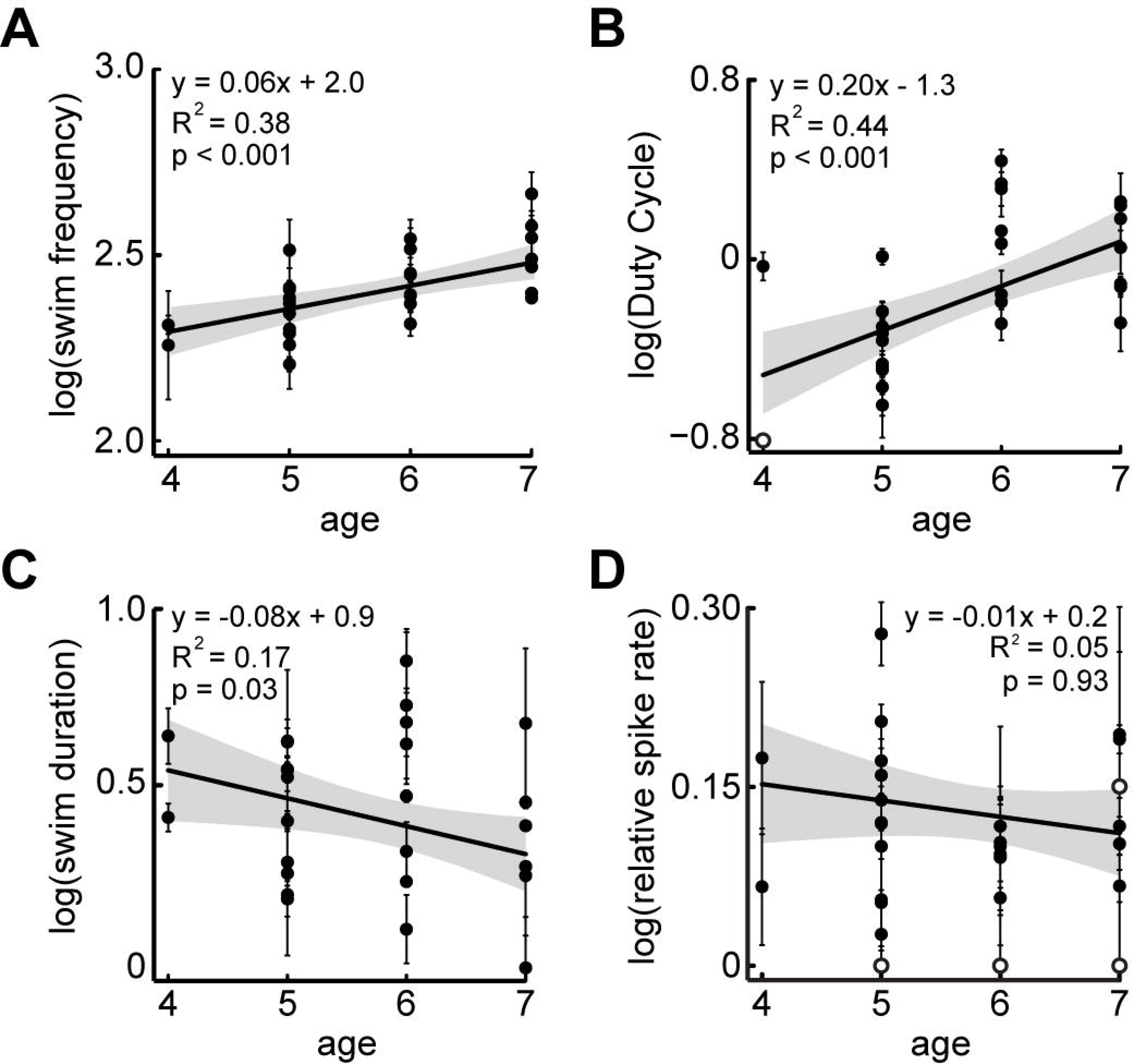
Effect of fish age on motor parameters and afferent inhibition. Swim frequency (**A**) and duty cycle (**B**) increases as larvae mature. Swim duration (**C**) decreases with age. **D.** The relative change in spike rate from pre-swim to swim intervals did not change across the ages investigated. All values represent mean ± SEM.

**Figure 6.**
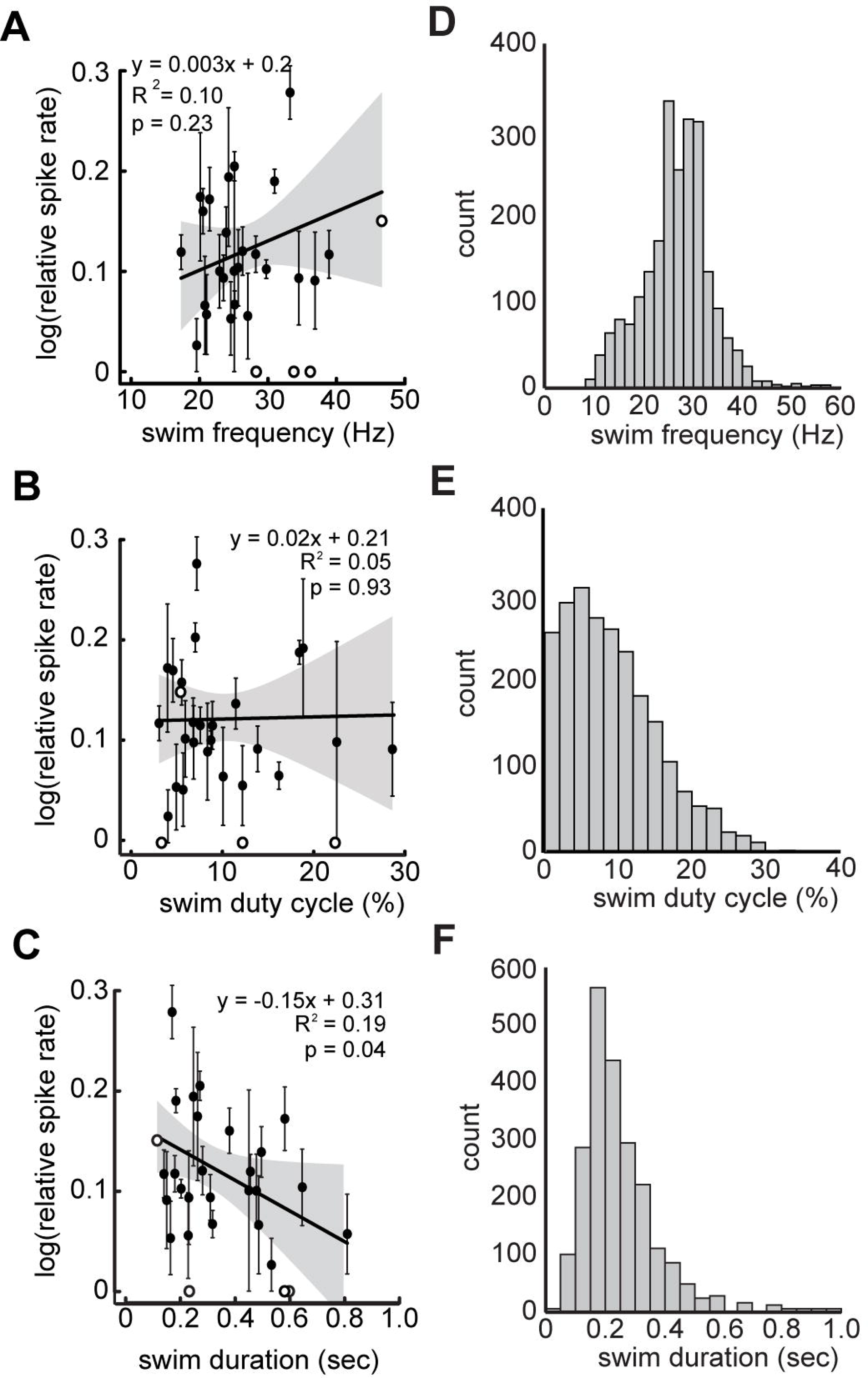
A longer swim duration leads to decreases in afferent spike rate. **A-B.** Swim frequency and swim duty cycle were not correlated to a decrease in afferent spike rate. **C.** Unlike frequency and duty cycle, swim duration was negatively correlated to afferent spike rate. The longer the swim duration, the higher the effect of inhibition. **D-F.** Histograms showing occurrences of swim frequency, duty cycle, and duration, respectively. All values represent mean ± SEM. Open dots represent individual with low statistical weight and we have omitted their error bars.

We characterized the reduction of afferent spike rate following a swim to define the refractory period and its contribution to post-swim spike dynamics. The time interval for post-swim spike rate to return to spontaneous levels was similar to the timing of when the first spike would be predicted to occur after a swim in control treatments when using a non-linear least squares model with a logistic function fit to spike rate (Figure 7 A-Bi). The same model indicated that the post-swim was relatively unchanged in ablation treatments. The density estimates of the time of the first spike after the swim was log-normally distributed in control treatments (Figure 7 Bii). In contrast, the time of the first spike in the ablation treatments could not be determined, resulting in a uniform distribution (Figure 7 Bii). The mean time to the first post-swim spike (0.149 ± 0.004 sec) was significantly later than the midpoint of the best-fit equation (t_mid_ CI = 0.105 – 0.119 sec), and within 95% confidence of the asymptotic spike rate (CI = 0.092 – 0.156 sec).

**Figure 7.**
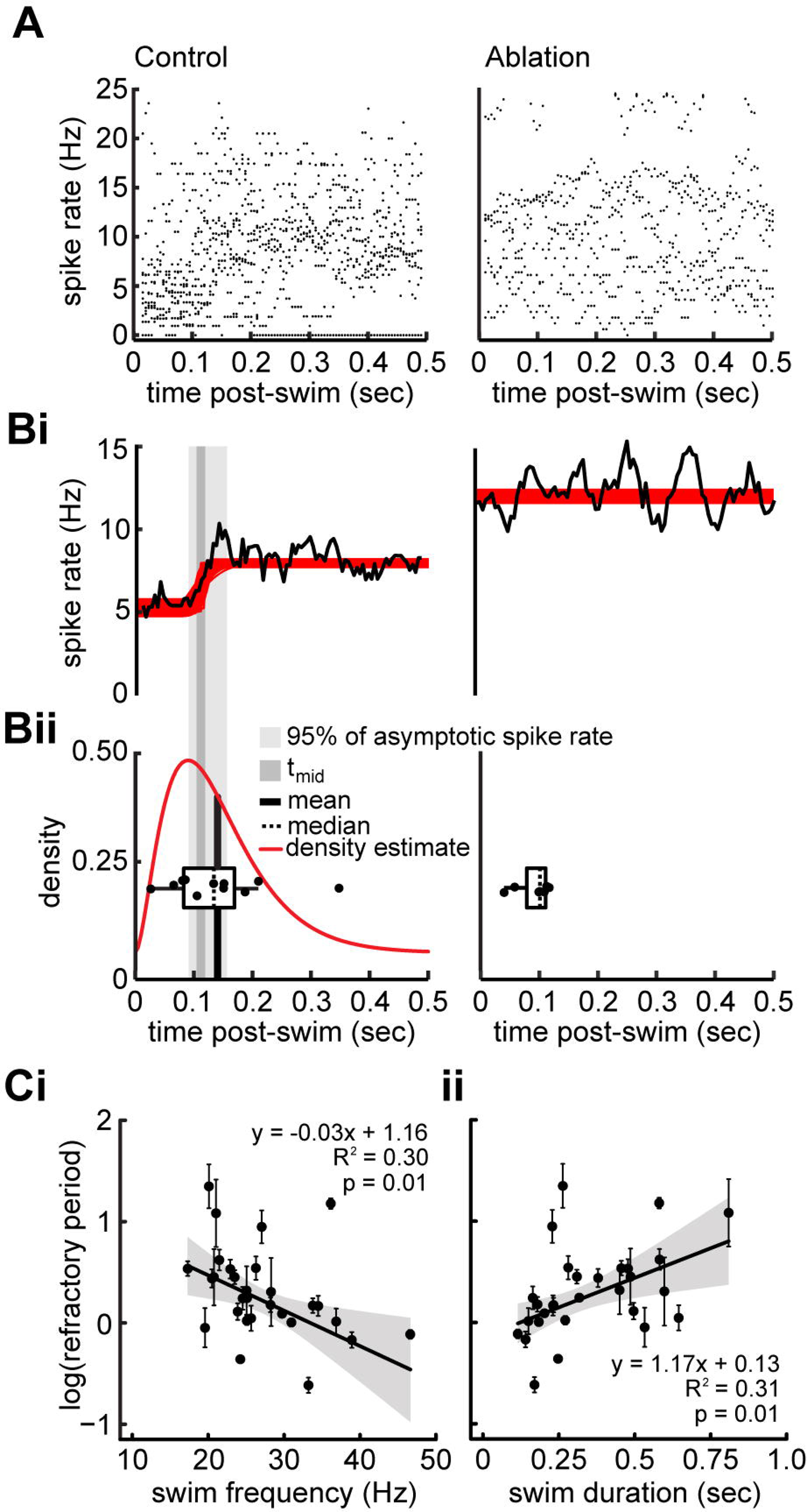
Spontaneous afferent spike rate remains suppressed after the offset of swimming. **A.** Raster plot of the instantaneous afferent spike rates within a 0.5 second window after the swim in both control (left) and ablation (right) larvae. **Bi.** Boot-strap simulation of spike rate (red line) and average spike rate (black line) reveal reduced afferent activity during the refractory period and a gradual return to intrinsic spontaneous spike rates in control treatments, and relatively consistent spike rates after the swim in ablation treatments. **Bii.** Box-plot of median refractory period (dashed line) for each individual overlaid atop the density estimate (red line) of the refractory period and the mean of the estimates (black bar). Both median and mean refractory periods are within the 95% confidence interval of the asymptotic spike rate for control treatments. There was no asymptote detected in ablation treatments. **C.** Linear regression between refractory period and swim frequency (i) and swim duration (ii). Each point represents the mean value for an individual ± SEM.

The mean value for the time to the first post-swim occurred within the confidence interval of 95% of the asymptotic spike rate, indicating that the timing of the first post-swim spike relative to the termination of the swim is a reliable predictor of refractory period duration (Figure 7 Bi-ii). Spike rate following ablation did not exhibit a quantifiable refractory period. Because the first post-swim spike time appears to coincide with the population-averaged refractory period, we use it to explore post-swim dynamics in control larvae. Refractory period duration was negatively correlated to the swimming frequency (r^2^ = 0.296, F_2,26_ = 5.467, p = 0.010; Figure 7 Ci), was not correlated with duty cycle (r^2^ = 0.088, F_2,26_ = 1.255, p = 0.998), and was positively correlated with swim duration (r^2^ = 0.315, F_2,26_ = 5.97, p = 0.007; Figure 7 Cii). Refractory period duration also increased as the level of inhibition increased (r^2^ = 0.391, F_2,26_ = 8.335, p = 0.001).

We quantified the swimming kinematics and glide duration of freely moving larvae to determine examine potential relationships between motor and sensory properties of the swim. Although refractory period duration was dependent on swimming frequency and duration, glide duration (0.49 ± 0.02 seconds) was independent of both tail-beat frequency (r^2^ < 0.001, F_1,94_ = 0.014, p = 0.905; Figure 8 B) and swim duration (r^2^ = 0.015, F_1,109_ = 1.69, p = 0.196; Figure 8 C). Consequently, the proportion of the glide affected by the refractory period was expected to decrease with faster tail-beat frequencies (r^2^=0.25, F_1,94_ = 31.35, p < 0.001; Figure 8 D) and increased with longer swim durations (r^2^ = 0.05, F_1,109_ = 6.09, p = 0.015). On average, the modeled refractory period accounts for the first 30.4% of the glide phase.

**Figure 8.**
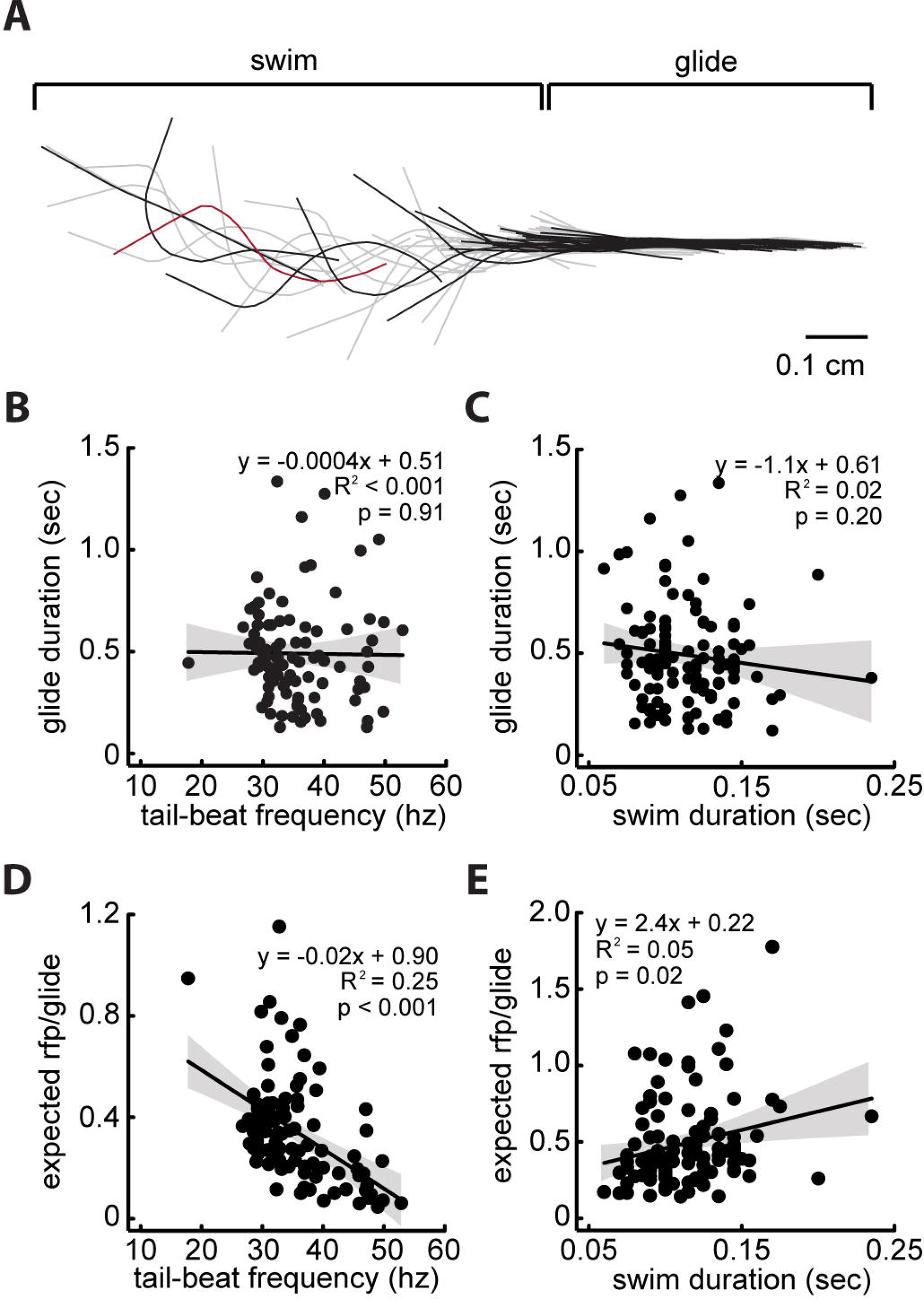
Tail-beat frequency and swim duration do not influence glide duration. **A.** Midline of zebrafish during swim and subsequent glide. Midline denoted in black every 4 ms. Linear regression between glide duration and tail-beat frequency (**B**) and swim duration (**C**). Linear regression between the proportion of the glide duration masked by its expected refractory period (rfp; calculated from linear model, see Figure 7 D) and tail-beat frequency (**D**) and swim duration (**E**).

## Discussion

When animals move their sensory receptors should, in theory, respond to a stimulus regardless of whether it is generated by the external environment or from the animal’s own movements. These movements can create sensory overload during locomotion and potentially masks stimuli important for survival, such as those generated by predators or prey. One solution would be for the animal to adjust its receptor sensitivity in anticipation of moving the body platform in which the sensory structures are embedded (von Holst and Mittelstaedt 1950; Sperry 1950; Crapse and Sommer 2009).

The spontaneous activity of zebrafish lateral line afferent neurons may be related to the maintenance of sensitivity. In other sensory systems, the degree of spontaneous firing is positively correlated with sensitivity to external stimuli (Manley and Robertson 1976, Kiang et al. 1965; Douglass et al. 1993; Yu et al. 2004). Here we describe the properties of a circuit that may regulate lateral line sensitivity during swimming motor commands (Feitl et al. 2010; Mensigner et al. 2019). We found that during 70% of swims there was a reduction in spontaneous afferent activity. This highlights the importance of, recording both afferent and motor activity together. Otherwise, spontaneous afferent activity will be underestimated due to the influence of swim-correlated inhibitory efferent activity (Harris and Milne 1966; Dawkins and Sewell 2004; Trapani and Nicolson 2011, Liao and Haehnel 2012; Haehnel-Taguchi et al. 2014; Song et al. 2018). This is particularly problematic when deciphering the underlying dynamics of afferents; intermittent swims can explain apparent state transition in afferent spike intervals (Song et al. 2018). The variability in spike rate response between afferent neurons (Levi et al. 2014; Haehnel-Taguchi et al. 2014) may also be influenced by efferent modulation. Additionally, we show that spike rate modulation is not limited to the swim period, but exhibits a nonlinear return of pre-swim spike rates after the swim ends. Characterizations of afferent activity should therefore also account for the delayed return of intrinsic spike properties after a swim.

Cholinergic efferents exert modulatory effects on the activity of the lateral line and other octavolateralis systems, such as the vestibular and auditory system. This has been demonstrated by galvanic stimulation to efferents, bath applied agonists, and genetic manipulation, all of which lead to a decrease in afferent activity (Russell 1971; Flock and Russell 1973; Russell and Roberts 1972; Vetter et al. 2007; Sewell and Starr 1991; Dawkins et al. 2007). These methods do not specifically target the efferent system or capture the intrinsic efferent activity. We note that while exogenous manipulations can elucidate the general function of physiological processes, they may be inadequate when tasked with revealing the nuanced *in vivo* dynamics inherent in endogenous systems. While traditional methods can document the general effect of the efferent system, it struggles to reveal its function in a natural context, where native circuit architecture and neurotransmitter levels are preserved. Using optogenetics we verified that cholinergic hindbrain efferents are synchronously active with spinal motor neurons during swimming (Bricaud et al. 2001; Chagnaud et al. 2015). We targeted and ablated these efferents and found that the magnitude of suppression was greatly decreased compared to controls. Residual suppression was expected because our method could not discriminate whether all efferents in the population were ablated. We found that efferents are necessary to inhibit afferent activity because when ablated there was no inhibition of afferent activity. However, efferents are not sufficient in compensating for the physiological heterogeneity of afferent neurons that contributes to variation in the level of inhibition. This is because even when efferents were intact, afferent activity was not always completely absent during swimming.

In order to survive, many animals must move across a wide range of speeds. In many aquatic animals an increased swimming frequency proportionally reduces afferent firing rate, likely due to the input of efferent neurons conveying this frequency information from active motor units (Russell 1971; Russell and Roberts 1972; Roberts and Russell 1972; Flock and Russell 1973; Hänzi et al. 2015; Chagnaud et al. 2015). In zebrafish, these speeds are achieved by increases in tail-beat frequency (Li et al. 2012), which are generated in a tonotopic manner by shifts in activity by distinct groups of interneurons and motor neurons in the spinal cord (Mendell 2005; McLean et al. 2007; McLean and Fetcho 2008). In paralyzed preparations, the tail-beat frequency is reflected in motor burst frequency (McLean et al. 2008). We examined variables of the motor command to better understand which ones might be copied by the efferent neurons. If motor burst frequency is the same as tail-beat frequency, which is the parameter commonly used to measure swim speed in zebrafish (Buss and Drapeau 2001; Liao and Fetcho 2008; McLean et al. 2008; Gabriel, et al. 2011; Severi et al. 2014), then we would expect to see that a relationship between motor burst frequency and afferent spike rate. However, we found no changes in afferent spike rate across swimming frequencies.

As larval zebrafish mature, swimming and other motor behaviors can change (Muller and van Leuwen 2004; McLean et al. 2008). In our hands, we found that average swim frequency increases with age, while swim duration decreases with age. We expected these age-related motor differences to be reflected in efferent modulation, but found no significant correlations. Despite the changes in motor commands with age the inhibitory effect remains constant. This suggests that the efferent system acts as a corollary discharge with limited temporal information from higher order motor units rather than an efference copy that encodes the timing from local motor units (Strakas et al. 2018). We investigated using a limited age range, but we suggest expanding to more mature fish to determine if the relationship between motor command and inhibition changes with development.

Swim duration had the highest correlation to afferent spike rate. Longer swim durations resulted in lower afferent spike frequencies, suggesting that information is filtered the periphery during extended swim bouts. Swim duration reflects an underlying mechanism of locomotor drive that corresponds to the strength of efferent inhibition. Our findings are consistent with auditory efferents that carry corollary discharges of vocalization motor commands in midshipman fish (*Porichthys notatus*). Auditory efferents have been shown to be adapted for the transmission of call duration information (Weeg et al. 2005; Chagnaud et al. 2011; Chagnaud and Bass 2013). In midshipman, there is evidence that the corollary discharge originates in the pre-pacemaker cells of the hindbrain which control gross timing (i.e. duration) of the motor command (Chagnaud et al. 2011; Chagnaud and Bass 2013), whereas control of motor command frequency occurs downstream of the corollary discharge. Additionally, weak encoding of swim frequency in the periphery may be due to persistent postsynaptic hair cell or afferent responses (e.g. neurotransmitter washout) in tadpole and toadfish (Highstein and Baker 1986; Boyle et al. 2009; Chagnaud et al. 2015).

We found that the reduction of afferent spike rate persisted beyond the offset of swimming revealing that the influence of efferent activity is not confined only to periods of active motor commands. This reduction was substantial, returning to intrinsic spontaneous spike rates after a well-defined refractory period. The refractory period is likely the product of either the interruption of the spontaneous spike pattern or a persistent hyperpolarizing effect of the efferents (Flock and Russell 1976; Art et al. 1982). This observation is supported by the fact that the refractory period was correlated to relative spike rate during swimming, swim frequency, and duration, suggesting that reduced spike rate is related to motor correlates within the efferent signal.

The refractory period has important implications for sensing in freely swimming larval fishes that employ intermittent burst-glide swimming patterns. The glide behavior following swim bouts may allow efficient sensing because the straightened body mitigates complex wake interactions that might interfere with responses to external stimuli (McHenry and Lauder 2005; Feitl et al. 2010; Tan et al. 2011). We found that the glide duration is not directly influenced by tail-beat frequency or swim duration, but can still be influenced by other factors such as could potentially be explained by differences in mechanical scaling (Muller et al. 2000; Lauder and McHenry 2005) or viscosity of the fluid environment at different depths (Anderson et al. 2001). Because the swim influences the refractory period and the duration of the glide phase is relatively constant across swimming parameters, on average, the refractory period accounts for 30% of the glide duration. The relationship between the refractory:glide duration and tail-beat frequency or swim duration reveals that fast, short swim bouts minimizes lateral line desensitization during the glide period. In light of our results, we must consider that gliding fish are at least partially desensitized to hydrodynamic signals, and this desensitization is influenced by the preceding swim. The refractory period may compensate for reafference generated by inertial passive movement, but it may also attenuate the lateral line during crucial opportunities for flow stimuli sampling. Therefore, the long-standing idea that intermittent or burst-and-glide swimming exists for sensory advantage is potentially inflated (Feitl et al. 2010; Tan et al. 2011). Neurophysiological mechanisms must be taken into consideration when attempting to interpret how movement strategies may influence the sensory capabilities of animals.

We still know relatively little about efferent modulation in freely-swimming fishes, where hydrodynamic stimuli from swimming can further impact afferent activity, but we now have a greater appreciation for the complex relationship between sensory and motor systems. In freely moving fish, afferent spike frequencies increase both during swimming and during feeding strikes (Palmer et al. 2005; Palmer et al. 2003; Ayali et al. 2009; Mensinger et al. 2019). This is consistent with our observation that efferent activity only partially suppresses spontaneous spike rates. To determine how efferents modulate evoked spike rates and ultimately relate to behavior, it is necessary to examine the magnitude of reafference in the absence of efferent activity, as we have done here. We have found that characterizing even basic properties of the afferent signal crucially depends on knowledge of the swim cycle. With the advent of new technologies and advances in both model systems and comparative approaches, neurophysiologists are primed to look deeper into the mechanisms of sensory feedback *in vivo* during locomotion and behavior.

## Acknowledgements

We gratefully acknowledge support from the National Institute of Health (DC010809), National Science Foundation (IOS1257150), and the Whitney Laboratory for Marine Biosciences to J.C.L.. We like to thank James Strother for generously sharing Tg(HUC:GCaMP6s) fish during preliminary experiments and thank Otar Akanyeti for programming assistance and comments on the manuscript.

## Author Contributions

E.T.L. and J.C.L conception and design of research; E.T.L. and D.A.S. performed experiments; E.T.L. and D.A.S. analyzed data; E.T.L, D.A.S., and J.C.L. interpreted results of experiments; E.T.L. prepared figures; E.T.L. drafted manuscript; E.T.L., D.A.S., and J.C.L. edited and revised manuscript; E.T.L., D.A.S., and J.C.L. approved final version of manuscript.

